# Reduced memory precision in older age is associated with functional and structural differences in the angular gyrus

**DOI:** 10.1101/2022.05.26.493542

**Authors:** S. M. Korkki, F. R. Richter, H. M. Gellersen, J. S. Simons

## Abstract

Decreased fidelity of mnemonic representations plays a critical role in age-related episodic memory deficits, yet the brain mechanisms underlying such reductions remain unclear. Using functional and structural neuroimaging, we examined how changes in two key nodes of the posterior-medial network, the hippocampus and the angular gyrus, might underpin loss of memory precision in older age. Healthy young and older adults completed a memory task that involved reconstructing object features on a continuous scale. Investigation of BOLD activity during retrieval revealed an age-related reduction in activity reflecting successful recovery of object features in the hippocampus, whereas trial-wise modulation of BOLD signal by graded memory precision was diminished in the angular gyrus. Grey matter volume of the angular gyrus further predicted individual differences in memory precision in older age, beyond likelihood of successful retrieval. These findings provide converging evidence for a role of functional and structural integrity of the angular gyrus in constraining the fidelity of episodic remembering in older age, yielding new insights into parietal contributions to age-related episodic memory decline.

## Introduction

Episodic memory, the kind of long-term memory that enables recollection of the spatio-temporal, perceptual, and emotional details that constitute a prior experience, exhibits marked declines with older age (Grady, 2012; Nyberg et al., 2012; Rönnlund et al., 2005). Memory changes typically seen in later life, including reduced recall specificity (Addis et al., 2008; Levine et al., 2002), deficits in mnemonic discrimination (Gellersen, Trelle, et al., 2021; Stark et al., 2013; Yassa et al., 2011) and increased vulnerability to interference (Pidgeon & Morcom, 2014; Wilson et al., 2018), suggest ageing to impact the level of detail and specificity with which information can be encoded into memory and subsequently retrieved. Indeed, recent behavioural work using continuous measures of long-term memory retrieval, where participants recreate aspects of studied stimuli (e.g., spatial location, colour) on an analogue scale, indicates decreased precision of memory representations to play a key role in age-related episodic memory deficits (Gellersen et al., 2023; Korkki et al., 2020; Nilakantan et al., 2018; Rhodes et al., 2020). Model-based analyses of the sources of participants’ retrieval errors in these types of tasks demonstrate that even when able to successfully retrieve details about a studied event from memory at a similar rate as younger adults, the fidelity of the details reconstructed is impoverished in older age (Korkki et al., 2020; Nilakantan et al., 2018; Rhodes et al., 2020). These age-related reductions in mnemonic precision appear in large part to result from differences in long-term memory retrieval (Korkki et al., 2020), however, the neural mechanisms underlying impoverished retrieval fidelity in older age remain unknown.

Neuroimaging findings from younger adults implicate the angular gyrus (AG), a ventrolateral parietal node of the posterior-medial network involved in episodic remembering (Ranganath & Ritchey, 2012; Ritchey & Cooper, 2020), as critical for supporting the fidelity of episodic recollection. AG activity generally increases with successful episodic retrieval (Rugg & Vilberg, 2013), and correlates positively with objective and subjective measures of memory detail and quality (Kuhl & Chun, 2014; Richter et al., 2016; Tibon et al., 2019; Vilberg & Rugg, 2007, 2009). When examining inter-regional differences in memory-related activation of the posterior-medial network, activity in the AG has been found to dissociate from that of the hippocampus (HC) (Richter et al., 2016). Specifically, while hippocampal activity was found to scale with the categorical success versus failure of object feature retrieval, activity in the AG predicted continuous variation in the precision with which participants successfully reconstructed object features from memory (Richter et al., 2016). These findings are consistent with theoretical accounts proposing the AG to play a role complementary to the HC during episodic retrieval (Moscovitch et al., 2016; Rugg & King, 2018; Sestieri et al., 2017; Simons et al., 2022; Wagner et al., 2005). While hippocampal pattern completion facilitates initial memory access and drives the cortical reinstatement of mnemonic content (Danker & Anderson, 2010; McClelland et al., 1995; Staresina et al., 2012), the AG is thought to support sustained maintenance and elaboration of the information reinstated in multiple modality-specific cortical regions (Humphreys et al., 2021; Rugg & King, 2018; Simons et al., 2022), necessary for more fine-grained judgements about specific details recovered from memory.

While many studies examining the neural basis of age-related episodic memory impairments have focused on the role of the HC and the surrounding medial temporal cortices (e.g., Carr et al., 2017; Reagh et al., 2018; Yassa et al., 2011), ageing also impacts functional and structural integrity of the wider cortico-hippocampal network (Andrews-Hanna et al., 2007; Reagh et al., 2020; Salami et al., 2014; Sun et al., 2016). Critically, age-related reductions in ventrolateral parietal activation during episodic memory retrieval have been commonly observed in prior studies (Wang & Cabeza, 2016), often co-occurring with decreases in hippocampal activity (Daselaar et al., 2006). Despite efforts to disentangle the relative contribution of these two regions to age-related episodic memory deficits, the specific behavioural consequences of parietal dysfunction in ageing remain unclear. Considering prior findings from younger adults (Richter et al., 2016), it is possible that age-related decreases in functional and structural integrity of the ventrolateral parietal cortex may underpin the loss of mnemonic precision observed in older age (Korkki et al., 2020; Nilakantan et al., 2018; Rhodes et al., 2020). Others have also emphasized a contribution of medial temporal regions to detailed episodic remembering (Moscovitch et al., 2016; Robin & Moscovitch, 2017; Yonelinas, 2013). For instance, the fidelity of spatial location retrieval has been shown to be diminished in patients with medial temporal lesions (Nilakantan et al., 2018), and to correlate with oscillatory activity in the hippocampus during episodic memory retrieval (Stevenson et al., 2018). Yet, the relative importance of these two regions in constraining memory fidelity in older age has not been directly evaluated in prior studies that have relied on categorical assessment of discrete retrieval outcomes (e.g., old/new, remember/know), which do not allow the mapping of brain function or structure to more graded variation in the resolution of mnemonic representations. As such, it is currently unresolved to what extent functional and structural changes in the hippocampus and/or the angular gyrus are responsible for age-related reductions in memory precision.

In the current study, we employed functional and structural magnetic resonance imaging (MRI) in combination with a continuous report paradigm to characterize the neural substrates of age-related decline in episodic memory precision. In the MRI scanner, healthy young and older adults encoded everyday objects that were presented in varying locations and colours on a scene background. At retrieval, participants were asked to recreate these features of the studied objects using a continuous, analogue scale, allowing for fine-grained assessment of memory retrieval. A further group of older adults underwent a structural MRI scan and completed the identical memory task outside of the scanner. Model-based analyses of participants’ retrieval errors allowed us to disentangle the neural substrates underpinning age-related alterations in the accessibility and fidelity of episodic memories, shedding new light on the functional consequences of cortico-hippocampal decreases for age-related episodic memory deficits.

## Methods

### Participants

Twenty-one younger (18-29 years old) and 53 healthy older adults (60-87 years old) took part in the current study. Two older participants failed to complete the full study and were therefore excluded. Functional and structural imaging data were acquired for all younger participants and for 21 of the older individuals, whereas 30 older individuals took part in a structural scan only and completed an identical memory task outside of the scanner. All participants were right-handed, native English-speakers, had normal or corrected-to-normal vision, no colour blindness, and no current or historical diagnosis of any neurological, psychiatric, or developmental disorder, or learning difficulty. Participants further indicated no current use of any psychoactive medication, and no medical or other contradictions to MRI scanning. Data from two older individuals was excluded from all analyses due to behavioural performance > 3 *SD*s from the group mean. For analyses of the fMRI data, one younger and one older adult were further excluded due to excessive movement in the scanner (> 4mm). Thus, the final sample sizes for the current analyses consisted of 20 younger and 19 older adults for the fMRI analyses and of 49 older adults for analyses concerning the structural data (see Table 1 for participant demographic information). All older participants scored within the healthy range (≥ 26, *M*: 28.51, *SD*: 1.34) on the Montreal Cognitive Assessment (MoCA) screening tool (Nasreddine et al., 2005). The young and older adults taking part in the fMRI experiment did not differ in terms of the number of years of formal education completed, *t*(37) = 0.44, *p* = .664, whereas the older adults had significantly higher scores on the Shipley Institute of Living Vocabulary Score (SILVS) (Zachary & Shipley, 1986) measure of crystallized intelligence, *t*(37) = 4.03, *p* < .001, *d* = 1.29, as typically observed in ageing studies (Verhaeghen, 2003). Moreover, the groups of older adults taking part in the fMRI scan and the structural part of the study only did not significantly differ in terms of age, education, MoCA or Shipley scores (*p*s > .118).

**Table 1.** Participant demographic information.

|  | Younger adults | Older adults |
| --- | --- | --- |
| N | 20 | 49 |
| Age | 22.15 (3.10) | 70.94 (5.32) |
| Gender (N) | 8 M, 12 F | 18 M, 31 F |
| Years of education | 16.90 (2.79) | 16.29 (4.75) |
| SILVS | 32.50 (3.91) | 36.98 (1.91) |
| MoCA | n/a | 28.51 (1.34) |

*Note.* M = males, F = females.

All participants were recruited via the University of Cambridge Memory Lab volunteer database, University of Cambridge Psychology Department Sona volunteer recruitment system (Sona Systems, Ltd), and community advertisements. Volunteers were reimbursed £30 for their participation and for any travel expenses incurred, and gave written informed consent in a manner approved by the Cambridge Psychology Research Ethics Committee.

### Materials

The memory task stimuli consisted of 180 images of everyday objects and 180 images of outdoor scenes that were randomly paired to form a set of 180 trial-unique encoding displays (size: 750 x 750 pixels) (see Figure 1). The location and colour of the object on each display were randomly sampled from circular parameter spaces (0-360°). All participants learned the same encoding displays.

**Figure 1.**
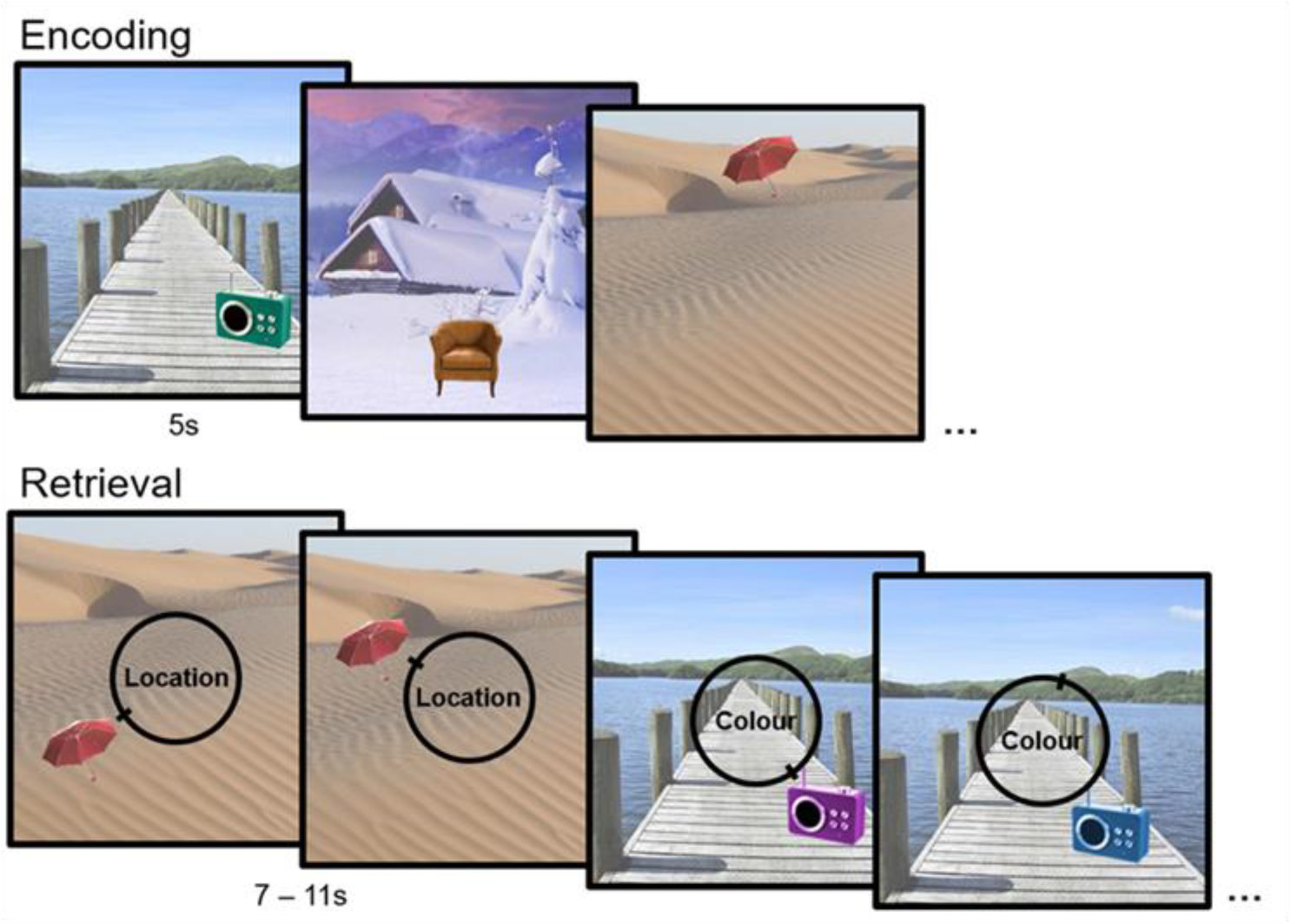
Memory task design. During encoding, participants viewed object-scene displays (stimulus duration: 5s). The location and colour of studied objects were randomly drawn from circular parameter spaces (0-360°). At test, participants were asked to recreate *either* the location *or* the colour of each studied object by moving a slider around a 360° continuous response dial (maximum response time: 11s). Retrieval error on each trial was measured as the angular difference between participants’ response value and the original encoded feature value.

### Design and procedure

The continuous report task has been described in detail in Korkki et al. (2021). A total of 9 study-test blocks were completed (see Figure 1). For the participants in the fMRI part of the study, each block was completed during one functional run (i.e., 9 functional runs in total, one study and one test phase per run). In each task block, participants first encoded 20 stimulus displays in a row (stimulus duration: 5s). After a 10s delay, they were asked to recreate *either* the location *or* the colour of each of the objects previously studied (one feature question per object, total of 20 retrieval trials per block). In the test phase, each object was presented on its original associated background. For location questions, the test object was presented in its original colour but in a location randomly drawn from the circular parameter space, whereas for colour questions the test object was presented in its original location but in a randomly chosen colour. Participants were instructed to recreate the location or the colour of the object as accurately as they could by moving a slider around a 360-degree response dial using their middle and index finger on a button box and confirmed their answer by pressing a third key on the button box with their thumb. The retrieval phase was self-paced but with a minimum trial length of 7s and a maximum response time of 11s. This maximum response time was based on older adults’ mean reaction time + 2 *SD*s in a fully self-paced pilot version of the task. If a participant failed to confirm their answer within the maximum allotted time, their last position on the response wheel was recorded as their answer for that trial.

In total, participants completed 90 location and 90 colour retrieval trials (10 trials of each type in each block). The feature questioned for each display was randomized, but kept constant across participants. Allocation of object-scene displays to task blocks and their study and test orders were randomized across participants with the constraint of no more than four sequential encoding or retrieval trials within the same feature condition. A fixation cross with a jittered duration ranging between 0.4s and 2.4s (mean: 1s) drawn from an approximate Poisson distribution was presented between each encoding and retrieval trial. For participants in the fMRI part of the study, a diffusion-weighted structural scan was acquired between functional runs five and six.

All participants completed instructions and a practice version of the memory task prior to the main task. For younger volunteers in the fMRI part of the study, the SILVS (Zachary & Shipley, 1986) was completed after the functional scan. For the older fMRI volunteers, the SILVS (Zachary & Shipley, 1986) and the MoCA (Nasreddine et al., 2005) were completed in a separate behavioural testing session. The older adults completing the structural scan only completed the continuous report task and the other behavioural measures on the same day as their structural scan.

### Analysis of behavioural performance

On each trial, retrieval error was calculated as the angular deviation between participants’ response and the original encoded target feature value (error range: 0 ± 180°). To examine the sources of memory error contributing to participants’ performance, we fitted a two-component probabilistic mixture model consisting of a target von Mises distribution and a circular uniform distribution (Bays et al., 2009; Zhang & Luck, 2008) to each participant’s retrieval error data using maximum likelihood estimation (code available at: https://www.paulbays.com/code/JV10/index.php). This model assumes that trial-wise variation in participants’ performance arises from two different components of memory error: variability, or noise, in successful retrieval of the target features from memory, and the presence of guess responses where memory retrieval has failed to bring any diagnostic information about the target to mind. Variability in successful memory retrieval is modelled by a von Mises distribution (circular equivalent of a Gaussian distribution) centred at the target feature value (i.e., mean retrieval error of zero), whereas the likelihood of guessing responses is captured by the probability of responses stemming from the circular uniform distribution. From this model, two parameters of memory performance can be estimated for all participants: the precision of memory retrieval, corresponding to the concentration parameter, *Kappa*, of the von Mises distribution, and the likelihood of successful memory retrieval, corresponding to the probability of responses stemming from the target von Mises over the uniform distribution (*pT* = 1 – *pU*).

Although alternative models can also capture the distribution of errors in continuous report tasks well (e.g., Bays, 2014; Schurgin et al., 2020; van den Berg et al., 2012), the selection of this model was motivated by prior work demonstrating differential effects of age on the two mixture-model components (Korkki et al., 2020; Rhodes et al., 2020), as well as differential neural correlates of these components in younger adults (Richter et al., 2016). Indeed, while agnostic to the specific mechanisms underpinning errors resembling variability in the accuracy of target retrieval and random guessing, we note that this model provides a good descriptive account of data generated in continuous report tasks (Bays & Taylor, 2020), and has been widely applied in both behavioural and neuroimaging studies of long-term memory retrieval (e.g., Brady et al., 2013; Cooper et al., 2017; Cooper & Ritchey, 2019; Richter et al., 2016; Stevenson et al., 2018; Sutterer & Awh, 2016). It is worth noting that a simpler one-parameter model, capturing recall errors as a noisy familiarity signal when taking the psychophysical similarity function of the stimulus space into account, has been proposed to account for distribution of retrieval errors in both working and long-term memory (Schurgin et al., 2020). However, assumptions of this model have been challenged (Tomić & Bays, 2022), and it is inconsistent with many existing models of hippocampal retrieval, which do not consider hippocampal function as a familiarity-like process (Norman, 2010; Norman & O’Reilly, 2003; Yonelinas et al., 2010). We further replicated our functional and structural neuroimaging analyses using model-free metrics of behaviour (i.e., raw retrieval error) and have reported these in the Supplementary material. No significant age differences in functional activity, or relationships between GM volume and memory performance, were observed when employing model-free metrics of memory performance, suggesting a benefit of the mixture modelling approach over analysis of raw error metrics for characterizing age-and performance-related variation in the current dataset. The two-component mixture model further fitted the behavioural data better than an alternative model consisting of a target von Mises distribution only for both young (mean Bayesian Information Criterion (BIC) for one-component von Mises model: 386.10; mean BIC for von Mises + uniform mixture model: 317.62) and older adults (mean BIC for one-component von Mises model: 483.66, mean BIC for von Mises + uniform mixture model: 450.19).

### MRI acquisition

MRI scanning was performed at the University of Cambridge Medical Research Council Cognition and Brain Sciences Unit using a 3T Siemens Tim Trio scanner (Siemens, Germany) with a 32-channel head coil. For each participant, a high-resolution whole brain anatomical image was acquired using a T1-weighted 3D magnetization prepared rapid gradient echo (MPRAGE) sequence (repetition time (TR): 2.25s, echo time (TE): 3ms, flip angle = 9°, field of view (FOV): 256 x 256 x 192mm, resolution: 1mm isotropic, GRAPPA acceleration factor 2). The functional data were acquired over 9 runs using a single-shot echoplanar imaging (EPI) sequence (TR: 2s, TE: 30ms, flip angle° = 78, FOV: 192 x 192mm, resolution: 3mm isotropic). Each functional volume consisted of 32 sequential oblique-axial slices (interslice gap: 0.75mm) acquired parallel to the anterior commissure – posterior commissure transverse plane. The mean number of volumes acquired per functional run was 167.39 (*SD*: 7.49) and did not significantly differ between the age groups (younger adults: 166.09, *SD*: 8.08; older adults: 168.75, *SD*: 6.77, *t*(37) = 1.11, *p* = .273). The scanning protocols also included additional imaging sequences not analysed here.

### fMRI preprocessing and analyses

Preprocessing and analysis of both the functional and structural images were performed with Statistical Parametric Mapping (SPM) 12 (https://www.fil.ion.ucl.ac.uk/spm/) implemented in MATLAB R2021b. The first five volumes of each functional run were discarded to allow for T1 equilibration. Any additional volumes acquired after each task block had finished were also discarded so that the last volume of each run corresponded to a time point of ∼2s after the last fixation cross for each participant. The functional images were spatially realigned to the mean image to correct for head motion and temporally interpolated to the middle slice to correct for differences in slice acquisition time. The anatomical image was coregistered to the mean EPI image, bias-corrected and segmented into different tissue classes (grey matter, GM; white matter, WM; cerebrospinal fluid, CSF). These segmentations were used to create a study-specific structural template image using the DARTEL (Diffeomorphic Anatomical Registration Through Exponentiated Lie Algebra) toolbox (Ashburner, 2007). The functional data was normalized to MNI space using DARTEL and spatially smoothed with an isotropic 8mm full-width at half-maximum (FWHF) Gaussian kernel.

To gain trial-specific estimates of the success and precision of memory retrieval for the fMRI analyses, the two-component mixture model (von Mises + uniform distribution) was fitted to all retrieval errors across all participants in the fMRI study (7020 trials in total) to calculate a cut-off point at which the probability of participants’ responses stemming from the target von Mises distribution was less than .05 and responses thus likely reflected guessing (c.f., Cooper et al., 2017; Korkki et al., 2021; Richter et al., 2016). This cut-off point of ± 59 degrees was then used to classify each retrieval trial as successful (absolute retrieval error ≤ 59 degrees) or unsuccessful (absolute retrieval error > 59 degrees). For successful retrieval trials, a trial-specific measure of memory precision was further calculated as 180 – participant’s absolute retrieval error on that trial so that higher values (smaller error) reflected higher precision (range: 121 – 180). A measure of memory precision was not considered for the unsuccessful trials as responses in this condition were approximately randomly distributed and thus were not expected to carry meaningful information about memory quality. Indeed, in a control analysis that included an additional parametric regressor reflecting trial-wise variation in memory error for trials classified as unsuccessful, we did not observe any significant associations between BOLD activity and memory error in either of the two ROIs, or across the whole brain (see Supplementary material).

First-level General Linear Model (GLM) for each participant contained four separate condition regressors corresponding to successful location retrieval, successful colour retrieval, unsuccessful retrieval, as well as encoding. As done in previous work (Cooper et al., 2017; Richter et al., 2016), unsuccessful trials were modelled across the two feature conditions, due to low numbers of guessing responses per feature condition for some participants. For successful retrieval trials, trial-specific estimates of memory precision were further included as parametric modulators comprising two additional regressors in the model. The precision parametric modulators were rescaled to range between 0 and 1 to facilitate the direct comparison of success and precision-related activity, and mean-centred for each participant. Neural activity corresponding to the regressors of interest was modelled with a boxcar function convolved with the canonical hemodynamic response function (HRF), with event duration corresponding to the duration of the retrieval display on the screen (i.e., duration of 7s if participant’s RT on that trial was under 7s, and duration equal to participant’s RT if the RT exceeded 7s). Six participant-specific movement parameters estimated during realignment (3 rigid-body translations, 3 rotations) were included as covariates in the first-level model to capture any residual movement-related artefacts. Due to the small number of guessing trials in each functional run, data from all functional runs were concatenated for each participant, and 9 constant block regressors included as additional covariates. Autocorrelation in the data was estimated with an AR(1) model and a temporal high pass filter with a 1/128 Hz cut-off was used to eliminate low frequency noise. First-level subject-specific parameter estimates were submitted to second-level random effects analyses.

Contrasts for the fMRI analyses focused on examining retrieval activity that varied with the success and precision of memory retrieval. To examine retrieval activity associated with the success of memory retrieval, we contrasted successful retrieval trials (absolute retrieval error ≤ 59 degrees) to trials where memory retrieval failed (absolute retrieval error > 59 degrees; *retrieval success effects*). To identify retrieval activity associated with the precision of memory retrieval, we examined positive associations between BOLD signal and trial-wise variation in memory error on trials classified as successful (i.e., positive linear relationship between BOLD signal and precision parametric modulator; *retrieval precision effects*).

### Preprocessing and analysis of structural images

For analysis of structural data, we used voxel-based morphometry (VBM) (Ashburner & Friston, 2000) to examine associations between GM volume and memory performance. Preprocessing of the structural images for the VBM analyses included segmentation of the anatomical images into GM, WM, and CSF. These segmentations were then used to create a structural template image using the DARTEL toolbox (Ashburner, 2007), and normalised to MNI space using DARTEL. The normalized GM images were spatially smoothed with an isotropic 8mm FWHF Gaussian kernel, with modulation applied to preserve the total amount of GM within each voxel. For each participant, total intracranial volume (TIV) was computed as the sum of total GM, WM, and CSF estimates.

Two separate GLMs were constructed to examine the relationship of GM volume to memory performance within the full older adult sample: one including model-derived estimates of the probability of successful retrieval *(pT*) as the covariate of interest, and the other including memory precision (*Κ*) as the covariate of interest. For both models, participant age, gender, education level, and TIV were included as covariates of no interest. When comparing the two groups of older adults who took part in the fMRI scan versus the structural scan only, we observed lower probability of successful memory retrieval in the group of participants who performed the memory task inside the MRI scanner (*M*: 0.63, *SD*: 0.10) than in those participants who performed the task outside of the scanner (*M*: 0.71, *SD*: 0.12), *t*(47) = 2.53, *p* = .015, *d* = 0.73, perhaps reflecting an impact of the MRI scan environment on memory performance (Gutchess & Park, 2006). Memory precision, on the other hand, did not significantly differ between these two groups of participants (in-scanner *M*: 7.78, *SD*: 3.01, outside of the scanner *M*: 7.91, *SD*: 3.20), *t*(47) = 0.14, *p* = .888. To account for the differences in retrieval success, testing environment was added as an additional covariate in all VBM analyses.

### Regions of interest

The functional and structural neuroimaging analyses focused on two key regions of the posterior medial memory network: the HC and the AG. The selection of these regions was motivated by prior evidence suggesting a degree of functional specialization during episodic retrieval (Richter et al., 2016), as well as common findings of age-related alterations in both regions (Leal & Yassa, 2015; Wang & Cabeza, 2016). Left-lateralized ROIs were created with the Automated Anatomical Labelling (AAL) atlas, given evidence for left-lateralization of episodic retrieval effects (Spaniol et al., 2009). Statistical significance within each anatomical ROI was assessed using small-volume correction with a peak-level familywise error (FWE) corrected threshold of *p* < .05, correcting for the number of voxels in each ROI. In addition to the ROI analyses, we performed exploratory whole brain analyses to identify any additional regions displaying age differences in memory-related retrieval activity, or an association between GM volume and memory performance, at a whole-brain corrected threshold of *p* < .05 FWE-corrected.

## Results

### Differences in memory performance between young and older adults in the fMRI experiment

We first examined behavioural differences in memory performance between the young (N = 20) and older (N = 19) individuals who took part in the fMRI part of the study. For each trial, retrieval error was calculated as the angular deviation between the participant’s response value and the original encoded feature value (response – target, error range: 0 ± 180°, see Figure 2A for distribution of retrieval errors in each age group). The older adults exhibited significantly higher mean absolute retrieval error (*M*: 43.65, *SD*: 8.29) than the younger adults (*M*: 30.43, *SD*: 15.04), *t*(37) = 3.38, *p* = .002, *d* = 1.08, indicating overall poorer memory performance in the older group. To distinguish whether such performance reductions may reflect age-related decreases in the likelihood of successfully retrieving information from memory, or in the precision of the retrieved information, we further fitted the two-component probabilistic mixture model (Bays et al., 2009; Zhang & Luck, 2008) to each individual participant’s retrieval error distribution (see Methods for details and Figure 2A for aggregate model fits). Examination of the model parameters indicated that both the probability of successful retrieval of object features (i.e., probability of responses stemming from a von Mises distribution centred at the target object feature, *pT*), *t*(37) = 2.30, *p* = .027, *d* = .74, as well as the precision with which these features were retrieved (i.e., variability in memory error when retrieval was successful, estimated as the concentration parameter, *Kappa*, of the target von Mises distribution), *t*(37) = 4.61, *p* < .001, *d* = 1.48, were reduced in older compared to younger adults (see Figure 2B). Thus, when compared to the younger adults, the older adults demonstrated both reduced success of memory retrieval and reduced mnemonic precision, allowing us to further probe the neural correlates of these behavioural decreases.

**Figure 2.**
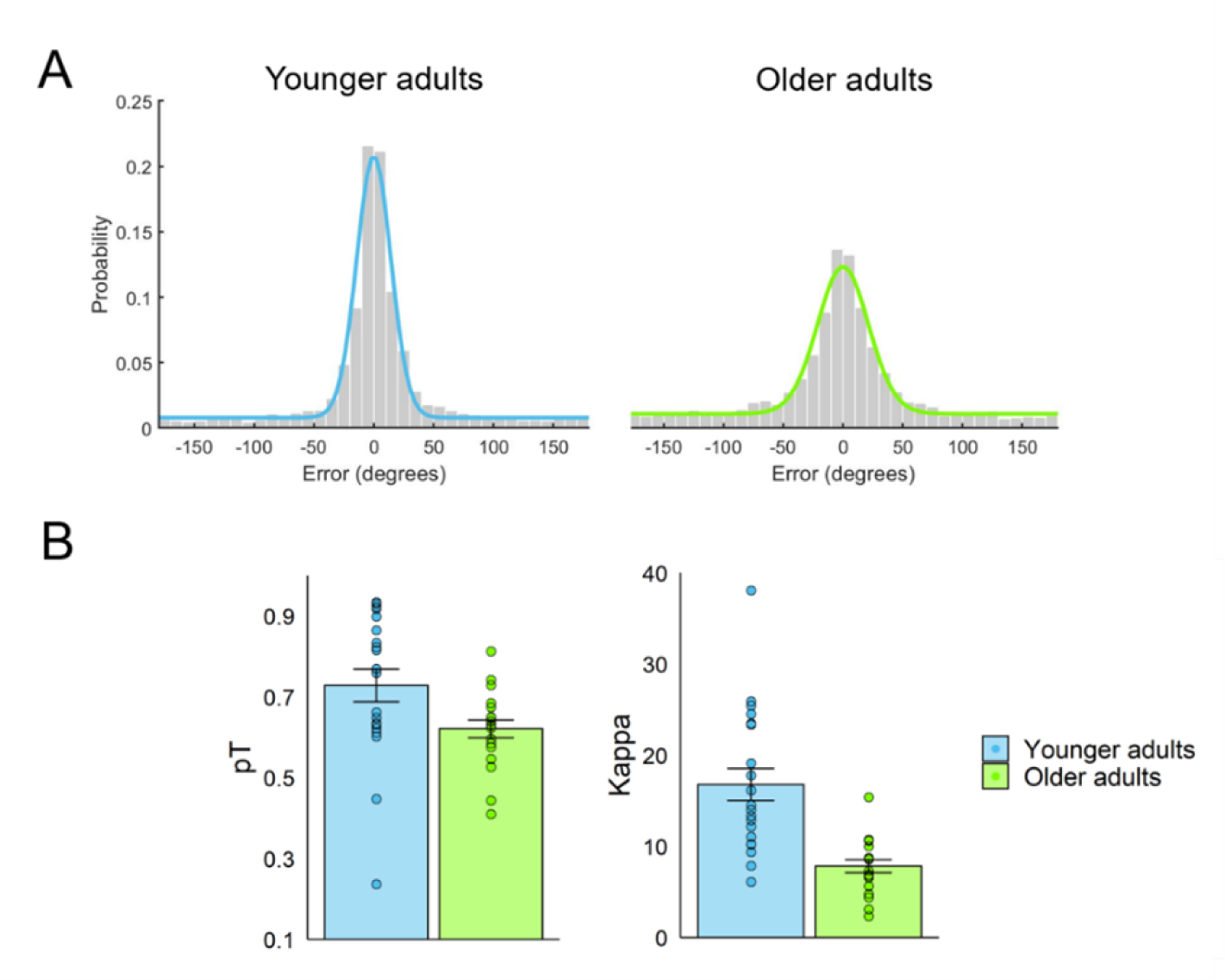
A) Distribution of retrieval errors across young (N = 20) and older participants (N = 19) in the fMRI experiment. Coloured lines indicate fits of the von Mises + uniform mixture model (Bays et al., 2009; Zhang & Luck, 2008), fitted to aggregate data in each age group for visualization. B) Mean model-estimated probability of successful memory retrieval (*pT*) and memory precision (*Kappa*) in each age group. Error bars display ± 1 standard error of the mean (SEM) and datapoints individual participant parameter estimates.

### Age differences in the hippocampus and the angular gyrus during memory retrieval

We next sought to examine age differences in memory-related activation of two key regions of the posterior-medial memory network, the HC and the AG, during retrieval. Across participants, we observed activity in both the HC, *t*(37) = 7.25, *p* < .001, peak: −24, −18, −12, and the AG, *t*(37) = 5.27, *p* = .001, peak: −39, −63, 21, to be significantly increased for successful compared to unsuccessful retrieval. Similarly, across participants, BOLD signal in the HC, *t*(37) = 5.63, *p* < .001, peak: −27, −15, −15, and the AG, *t*(37) = 6.13, *p* < .001, peak: - 54, −66, 33, correlated with trial-wise variation in mnemonic precision. Examining age-related differences in BOLD activity associated with these two aspects of memory retrieval, we observed significant age-related reductions in retrieval success effects in the HC, *t*(37) = 3.79, *p* = .024, peak = −18, −6, −12. Although increased hippocampal activity was detected for successful in comparison to unsuccessful trials in both younger, *t*(19) = 6.06, *p* = .001, peak = −30, −15, −12, and older adults *t*(18) = 4.37, *p* = .021, peak: −24, −18, −12, this effect was reduced significantly in older age (see Figure 3A). A non-significant trend in the same direction was observed in the AG, *t*(37) = 3.23, *p* = .093, peak = −39, −60, 21, with both young, *t*(19) = 5.48, *p* = .003, peak: −39, −63, 21, and older adults, *t*(18) = 4.47, *p* = .020, peak: −42, −54, 24, nevertheless demonstrating significant retrieval success effects in the AG also.

**Figure 3.**
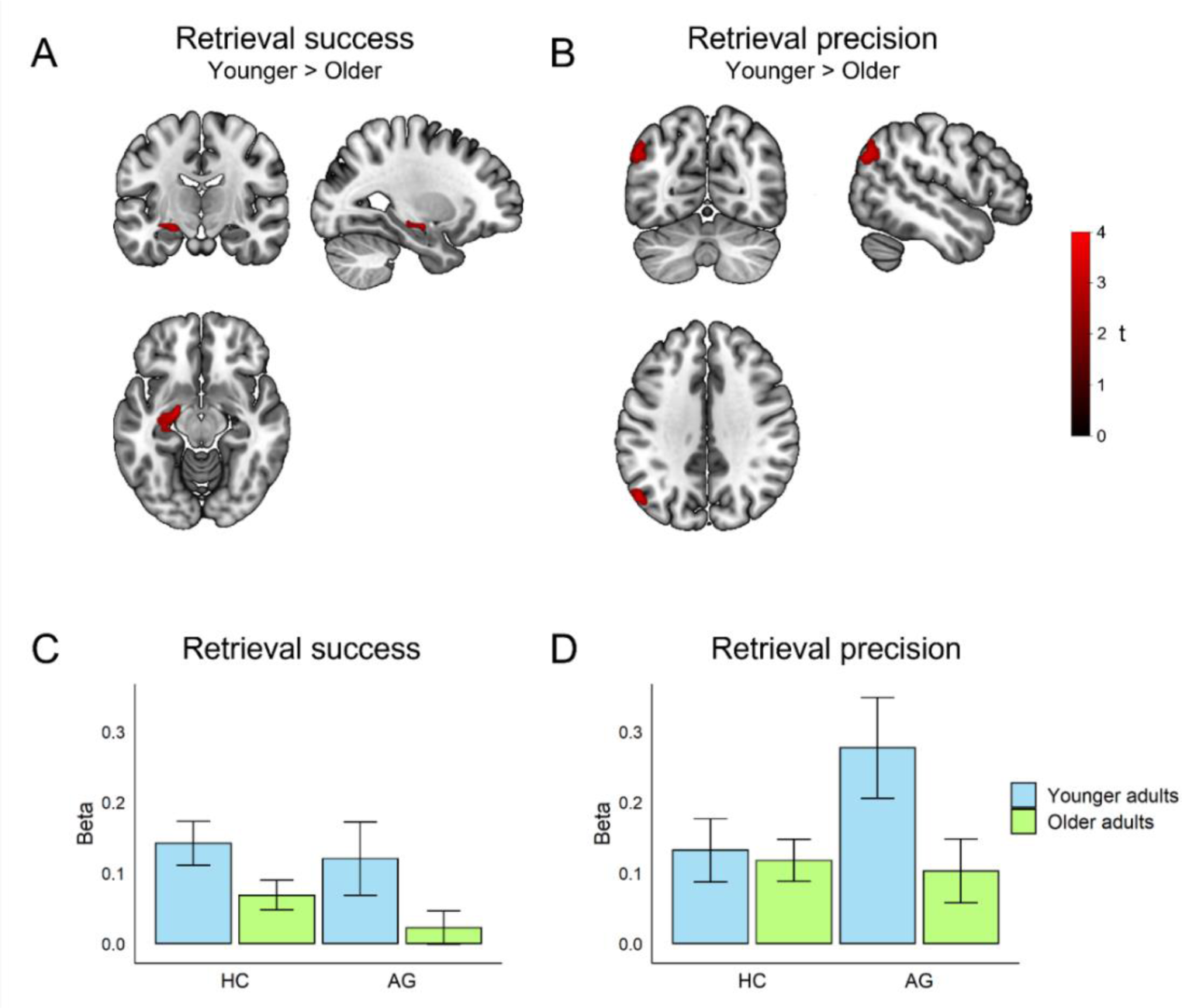
Age-related reductions (younger adults > older adults) in BOLD signal associated with the (A) success and (B) precision of memory retrieval within the (A) hippocampal and (B) angular gyrus ROI, displayed at an uncorrected threshold *p* < .01 for visualization. Mean beta-values for the (C) retrieval success and (D) retrieval precision effects in each age group and anatomical ROI. Error bars display ± 1 SEM.

Conversely, significant age-related reductions in retrieval precision effects were detected in the AG, *t*(37) = 3.67, *p* = .033, peak = −48, −69, 36 (see Figure 3B). Whereas in younger adults AG activity was significantly modulated by the precision of memory retrieval, *t*(19) = 5.50, *p* = .003, peak: −54, −66, 33, no significant precision effects were detected in the older adult group alone *(p*s > .152). Hippocampal activity associated with memory precision, on the other hand, did not significantly differ between the age groups (*p*s > .374). Indeed, significant modulation of hippocampal BOLD signal by memory precision was observed in both young, *t*(19) = 4.78, *p* = .008, peak = −27, −18, −18, and older adults, *t*(18) = 4.68, *p* = .012, peak = −30, −27, −15. No additional age differences in retrieval success or retrieval precision effects survived a whole-brain corrected threshold in exploratory whole brain analyses (*p*s > .077).

Given prior evidence for functional specialization of activity associated with the success and precision of episodic memory retrieval (Richter et al., 2016), we further extracted the mean beta-values corresponding to the retrieval success and retrieval precision effects from the anatomical HC and AG ROI, and subjected them to a 2 (region) x 2 (measure) x 2 (age group) ANOVA (see Figure 3C and 3D). This analysis indicated a significant 2-way interaction between region and measure, *F*(1,37) = 5.41, *p* = .026, *η_p_ ^2^* = .13. Across participants, we observed significantly greater retrieval precision than retrieval success effects in the AG, *t*(38) = 3.12, *p* = .003, *d* = .50, whereas activity associated with the success and precision of memory retrieval did not significantly differ in the hippocampal ROI (*p* = .546). There was no further 3-way interaction between age group, region, and measure (*p* = .120). However, a trend toward a 2-way interaction between age group and memory measure was observed for precision-related activity, *F*(1,37) = 3.57, *p* = .067, *η_p_ ^2^* = .09, which demonstrated marginally greater age-related differences in the AG when compared to the HC. The 2-way interaction between age group and memory measure was non-significant for retrieval success related activity (*p* = .348).

### Relationship between variation in local grey matter volume and memory performance in older age

Examining the full sample of older individuals who took part in the study (*N* = 49; memory task completed in-scanner for 20 individuals and outside of the scanner for 29 individuals), we further investigated whether individual differences in GM volume of the HC and the AG were related to variation in episodic memory performance. Controlling for sex, education, and testing environment, we observed a significant positive association between age and mean absolute retrieval error on the continuous report task, *r* = .33, *p* = .027, indicating that in addition to detecting differences between young and older adults, the task was sensitive to age-related variation in memory performance within the older adult sample. Examining variation in the model-derived estimates of memory performance, we observed a significant negative correlation between age and memory precision, *r* = -.49, *p* < .001 (Figure 4B), whereas no significant association between age and retrieval success was detected, *r* = -.08, *p* = .619 (Figure 4A). Indeed, the correlation between age and memory precision was significantly stronger than the association between age and the probability of successful retrieval, *z* = 2.36, *p* = .018.

**Figure 4.**
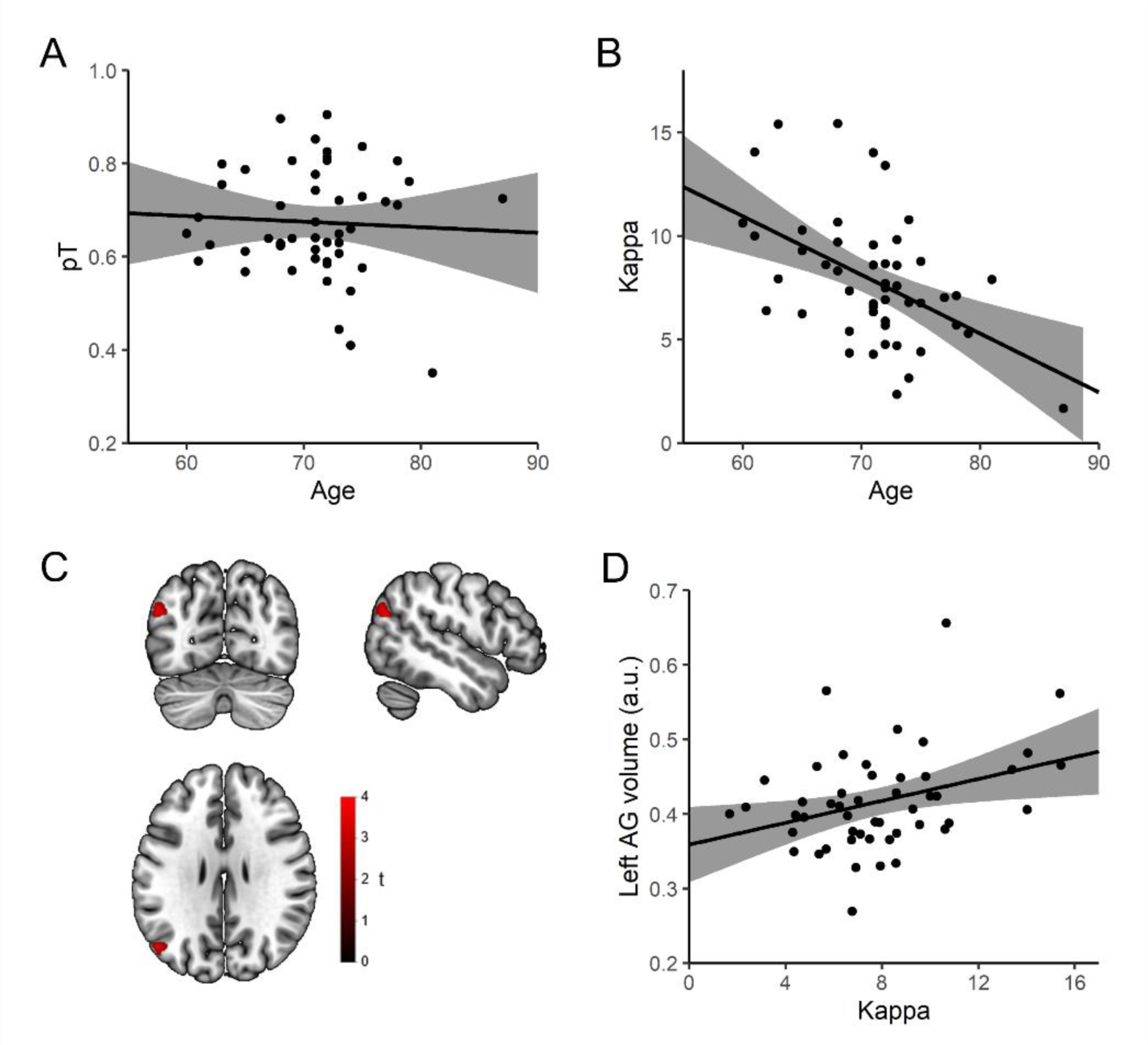
Relationship between age and the model-derived estimates of A) retrieval success and B) retrieval precision in the full older adult sample (N = 49). C) Angular gyrus region displaying an association between grey matter (GM) volume and memory precision in older age, visualized at an uncorrected threshold of *p* < .01. D) Relationship between angular gyrus GM volume and memory precision (GM volume extracted from the cluster showing a relationship to memory precision at *p* < .001 uncorrected).

Moreover, VBM (Ashburner & Friston, 2000) analyses indicated a significant positive association between AG GM volume and individual differences in memory precision, *t*(42) = 3.74, *p* = .026, peak: −46, −68, 24 (Figure 4C and 4D). In contrast, HC volume was not significantly associated with memory precision or the probability of successful memory retrieval within the full older adult sample (*p*s > .133). No significant association between AG GM volume and the probability of successful memory retrieval was detected (*p*s > .722). Indeed, when including probability of successful memory retrieval as an additional covariate in the model, we still observed a significant association between AG volume and memory precision, *t*(41) = 3.75, *p* = .027, peak = −46, −68, 24, suggesting specificity of this relationship to mnemonic precision. Exploratory whole-brain analyses did not reveal any further regions where variation in GM volume was associated with either the precision (*p*s > .185) or the success (*p*s > .789) of memory retrieval, although there was a trend toward a positive association between GM volume in the inferior temporal gyrus and memory precision, *t*(42) = 5.03, *p* = .061, peak: 50, −57, −9.

## Discussion

Here, we examined the contribution of two key regions of the posterior-medial memory network, the HC and the AG, to age-related episodic memory reductions. Employing continuous measures of long-term memory retrieval in combination with model-based analyses of participants’ retrieval errors, we observed both the probability of successful memory retrieval and mnemonic precision to be reduced in older compared to younger adults. Of these two measures, memory precision, but not the likelihood of successful retrieval, was also negatively associated with age within a larger older adult sample, indicating sensitivity of this measure for detecting age-related variation in memory performance. FMRI analyses of memory-related activation during retrieval revealed an age-related reduction in hippocampal activity reflecting the successful recovery of object features from memory, whereas activity associated with graded variation in memory precision was reduced in older age in the AG. Converging with the functional differences, AG GM volume also predicted individual differences in mnemonic precision in older age, beyond any relationship shared with successful memory retrieval. These findings highlight a contribution of declining functional and structural integrity of the AG to age-related loss of memory precision, shedding new light on cortico-hippocampal contributions to age-related episodic memory deficits.

Across participants in the fMRI study, we observed BOLD signal in both the HC and the AG to increase for trials that were associated with successful retrieval of the target object feature when compared to trials that reflected guessing. Aligning with previous work emphasizing a contribution of hippocampal dysfunction to age-related memory impairments (Hou et al., 2020; Leal & Yassa, 2015; Persson et al., 2012; Pudas et al., 2013; Trelle et al., 2020), we observed older age to be associated with significantly reduced retrieval success related activity in the hippocampus. This finding is consistent with prior studies reporting decreased categorical hippocampal recollection effects in older age (Cansino et al., 2015; Daselaar et al., 2006). Variation in hippocampal signal reflecting successful memory retrieval has previously been shown to predict both cross-sectional and longitudinal differences in episodic memory performance in later life (de Chastelaine et al., 2016; Hou et al., 2020), highlighting the functional significance of these decreases for memory function in older age. The hippocampal reductions observed here may in part reflect age-related deterioration of hippocampal pattern completion, a mechanism by which the HC enables the recovery and reinstatement of a complete encoded memory representation from a partial or noisy cue (McClelland et al., 1995; Norman & O’Reilly, 2003). This interpretation is consistent with previous reports of decreases in both behavioural and neural indices of pattern completion in ageing (Paleja & Spaniol, 2013; Trelle et al., 2020; Vieweg et al., 2015). Although age-related differences in retrieval success effects did not reach significance in the AG, we note that a trend-level reduction in BOLD signal reflecting successful memory retrieval was evident in this region also. As such, an additional contribution of extrahippocampal processes to reduced memory accessibility in older age cannot be excluded based on the current data. Indeed, given emerging insights about the temporal sequence of hippocampal and parietal retrieval operations, with hippocampal signal reflecting successful retrieval preceding that of the posterior parietal cortex (Martín-Buro et al., 2020; Treder et al., 2021), it is possible such a reduction could reflect impoverished input from the HC, which initiates memory retrieval, to the rest of the posterior-medial network in ageing.

Across participants, we also observed BOLD signal in both the HC and the AG to be positively associated with the graded precision with which participants reconstructed object features from memory. In contrast to retrieval success, we did not observe any significant age-related reductions in memory precision related activity in the HC, where trial-wise variation in BOLD signal during successful memory retrieval was positively associated with variation in the degree of memory error in both young and older adults. Instead, the older group displayed diminished precision-related activation of the AG, where significant modulation of BOLD signal by memory precision was limited to the younger adults. This finding suggests that the older adults were less able to flexibly modulate trial-wise activity in the AG to support detailed episodic remembering, consistent with recent evidence indicating diminished modulation of parietal event-related potentials by memory specificity in older age (Horne et al., 2020, but see Murray et al., 2019), as well as observations of age-related dysregulation of default-mode network (DMN) activity in response to cognitive demands more generally (Grady et al., 2006; Persson et al., 2007; Sambataro et al., 2010). The age-related differences in the AG were predicted by evidence from younger adults that emphasize a contribution of this region to qualitative aspects of episodic remembering (Rugg & King, 2018; Simons et al., 2022). Specifically, neuroimaging studies in younger adults have observed retrieval activity in the AG to correlate with the precision (Richter et al., 2016), detail-richness (Vilberg & Rugg, 2007, 2009), and vividness (Bonnici et al., 2016; Kuhl & Chun, 2014; Tibon et al., 2019) of episodic remembering, with evidence for a causal role further provided by studies demonstrating transcranial magnetic stimulation of the AG to impact subjective and some objective aspects of memory quality (Nilakantan et al., 2017; Thakral et al., 2017; Yazar et al., 2014; Zou & Kwok, 2022). Indeed, these findings align with patient evidence indicating lateral parietal lesions to be associated with subtle episodic memory deficits and altered subjective experience of remembering but not causing full-blown amnesia (Berryhill et al., 2007; Simons et al., 2010).

Although the ventrolateral parietal cortex consistently activates during episodic recollection (Rugg & Vilberg, 2013), its specific role in episodic memory has been debated (e.g., Cabeza et al., 2012; Rugg & King, 2018; Shimamura, 2011; Simons et al., 2022; Wagner et al., 2005). The current findings implicating the AG in age-related reductions in memory precision align with multiple lines of evidence emphasizing AG involvement in online representation of retrieved information (reviewed in Humphreys et al., 2021; Rugg & King, 2018; Sestieri et al., 2017). Following successful hippocampal pattern completion, which drives the initial cortical reinstatement of retrieved content (McClelland et al., 1995; Staresina et al., 2012), the AG is thought to facilitate the sustained maintenance and elaboration of event details (Ritchey & Cooper, 2020; Rugg & King, 2018), integrating information from multiple modality-specific cortical sites (Bonnici et al., 2016; Shimamura, 2011; Simons et al., 2022). This account is supported by evidence indicating multi-voxel neural representations in the AG during retrieval to code for individual memories (Bonnici et al., 2016; Kuhl & Chun, 2014) as well as for specific mnemonic features (Favila et al., 2018; Lee & Kuhl, 2016). The involvement of the AG in representing the contents of memory retrieval may reflect a more domain-general role of this region in facilitating the online buffering of multi-modal, spatio-temporally extended, representations (Humphreys et al., 2021). Indeed, this may be a critical computational role of the lateral parietal cortex at large, with graded differences in functional and structural connectivity with the rest of the brain (e.g., Caspers et al., 2011; Uddin et al., 2010; Wang et al., 2012) manifesting as involvement of parietal subdivisions in different behavioural functions (Humphreys & Tibon, 2023). While more dorsal parietal regions connect with frontal regions involved in cognitive control (Sestieri et al., 2017; Yeo et al., 2011), ventral parietal regions exhibit connectivity with medial temporal and DMN regions involved in episodic remembering (Humphreys & Tibon, 2023; Sestieri et al., 2017). Within the ventrolateral parietal cortex further anterior-posterior variation in functional and structural connectivity exists, with connectivity to medial temporal regions appearing strongest for the mid-ventrolateral parietal cortex (Humphreys et al., 2022). Our current results suggest that functional alterations in the AG in older age may compromise this online buffering of bound, multi-modal, event representations and the evaluation of specific aspects of these representations for precise mnemonic judgements. In addition to local alterations in the AG, it is possible that the age-related reductions observed here may in part reflect impoverished inputs to the AG from the hippocampus (Ritchey & Cooper, 2020; Robin & Moscovitch, 2017), or from modality-specific cortical regions involved in representation of particular event features (Danker & Anderson, 2010; Favila et al., 2020).

Converging with findings from the functional neuroimaging analyses, we further observed individual differences in AG GM volume to be positively associated with memory precision in older age. Critically, this relationship persisted after controlling for differences in the probability of successful memory retrieval, highlighting a specific contribution of structural integrity of the AG to mnemonic fidelity in older age. In addition to displaying functional alterations (e.g., Andrews-Hanna et al., 2007; Persson et al., 2014; Reagh et al., 2020), heteromodal cortical regions, including the ventrolateral parietal cortex, are particularly sensitive to age-related GM loss (Douaud et al., 2014; Fjell et al., 2014; McGinnis et al., 2011), with maintenance of GM integrity of these regions being associated with superior memory performance in older age (Sun et al., 2016). These findings suggest that the relationship between AG volume and memory precision observed here may in part reflect the degree of age-related structural deterioration of this region, although longitudinal data are required to fully evaluate this interpretation. Despite the functional age differences observed in the HC, we did not observe any significant associations between hippocampal volume and the success, or precision, of memory retrieval. Given the small size and structural complexity of the HC, and the contribution of different hippocampal subfields to distinct mnemonic processes (Hunsaker & Kesner, 2013), more fine-grained assessment of structural integrity of this region may be required to detect relationships with behaviour. Overall, the functional and structural neuroimaging results provided consistent support for a role of decreased integrity of the angular gyrus in loss of episodic memory precision in older age.

Although both the probability of successful memory retrieval and mnemonic precision were impaired in the older relative to the younger group, of these two measures only mnemonic precision displayed a significant negative association with age in the full older adult sample. The association between age and memory precision was stronger than any relationship between age and probability of successful retrieval, highlighting the potential benefit of this measure for detecting early age-related memory deficits. In addition to the functional and structural alterations observed with healthy ageing, the AG, along with rest of the DMN, is particularly vulnerable to early accumulation of Alzheimer’s pathology. Amyloid-β is found to accumulate early in regions of the DMN (Grothe et al., 2017; Palmqvist et al., 2017), contributing to functional alterations within this network (Ingala et al., 2021; Mormino et al., 2011; Palmqvist et al., 2017). An important avenue for future research to assess is whether continuous measures of long-term memory retrieval, such as the task used here, could be advantageous for detecting early signs of subtle pathological memory changes in ageing. Evidence from working memory assessment suggests continuous report paradigms to be sensitive to genetic risk of developing dementia (Lu et al., 2021; Pavisic et al., 2021; Zokaei et al., 2021), however the extent to which this link might extend to long-term memory remains unclear (Gellersen, Coughlan, et al., 2021).

While not afforded by the current task design, future work should also address whether the age-related differences in univariate activity observed here are coupled with reduced fidelity of multi-voxel neural representations of specific event features in the AG (Favila et al., 2018; Lee & Kuhl, 2016). Indeed, prior work has indicated ageing to be associated with decreased fidelity of cortical representations of retrieved content (Folville et al., 2020; St-Laurent et al., 2014; Trelle et al., 2020). Notably, reduced specificity of neural reinstatement in the AG was recently found to predict variation in episodic memory performance in older age beyond hippocampal retrieval activity in a large sample (Trelle et al., 2020), consistent with the idea that these two regions may contribute to distinct variation in memory performance in older age. However, we note that the current analyses provided only marginal evidence for an interaction between brain region and age group for precision related activity and no evidence for an interaction between brain region and age group for retrieval success related activity. As such, the regional specificity of age-related differences in activity associated with these two aspects of memory retrieval remains unclear based on the current data.

Despite common observations of age-related reductions in ventrolateral parietal activation during episodic retrieval (Wang & Cabeza, 2016), the functional significance of such reductions for memory deficits in older age has remained elusive. Here, using continuous report measures of long-term memory retrieval, we provide novel converging evidence for a contribution of functional and structural integrity of the AG to loss of memory precision in older age. Specifically, we observed older age to be associated with diminished modulation of AG retrieval activity by memory precision, with GM volume of this region further predicting variation in the fidelity of episodic remembering in later life. Further research is needed to explore the potential for memory precision measures to assist early detection of pathological age-related changes within the posterior-medial memory network.

## Acknowledgements

This study was funded by BBSRC grant BB/L02263X/1 and James S. McDonnell Foundation Scholar Award #220020333, and was carried out within the University of Cambridge Behavioural and Clinical Neuroscience Institute, funded by a joint award from the Medical Research Council and the Wellcome Trust. For the purpose of open access, the authors have applied a Creative Commons Attribution (CC BY) licence to any Author Accepted Manuscript version arising. We would like to thank the staff of the MRC Cognition and Brain Sciences Unit MRI facility for scanning assistance.

