## Supplementary material for "Reduced memory precision in older age is associated with functional and structural differences in the angular gyrus"

### *Retrieval activity and trial-wise variation in memory error for trials classified as unsuccessful*

As retrieval errors for trials classified as unsuccessful (i.e., absolute retrieval error > 59°) were approximately randomly distributed across the circular space (see main manuscript Figure 2A), we did not expect variation in memory error across these trials to be meaningfully associated with trial-wise variation in BOLD signal. In a control analysis, an additional parametric regressor reflecting trial-wise variation in memory error was included for trials classified as unsuccessful (180° – retrieval error, range: 0 – 120°). This analysis indicated no significant associations between memory error and BOLD signal in either the hippocampus (*p*s > .501), or the angular gyrus (*p*s > .906). Moreover, no significant associations between BOLD signal and variation in memory error for trials classified as unsuccessful were detected at a whole-brain corrected threshold either (*p*s > .901). These findings thus support the notion that variation in memory error on these trials was driven by guessing.

### *Retrieval activity and trial-wise variation in memory error collapsing across all trials*

We further examined whether any age differences in retrieval activity would be observed if examining associations between BOLD activity and trial-wise variation in memory error across all trials, without assuming a model-based distinction between successful and unsuccessful retrieval. For this purpose, an additional General Linear Model was constructed, modelling all location and colour retrieval trials with one regressor each and including a parametric modulator capturing trial-wise variation in memory error for each of these conditions. Although BOLD signal in both the hippocampus, $t(37) = 7.84$, $p < .001$, peak: -27, -15, -12, and the angular gyrus, $t(37) = 4.83$, $p = .002$, peak: -39, -57, 30, correlated with trial-wise variation in absolute error when collapsing across all retrieval trials, no significant age differences were observed in either the hippocampus ($p$s > .208) or the angular gyrus ($p$s > .093) in this analysis.

### *Grey matter volume and individual differences in mean retrieval error*

Additionally, we examined whether grey matter volume in the hippocampus or the angular gyrus was associated with individual differences in mean absolute retrieval error within the full older adult sample. A voxel-based morphometry analysis including mean absolute retrieval error as the covariate of interest did not indicate any significant associations between mean absolute error and grey matter volume in either the angular gyrus, ($p$s > .150) or the hippocampus ($p$s > .742).
